# Label-free acoustic and optical microscopy of live tumor spheroids in hydrogel for high-throughput 3D In-vitro drug screening

**DOI:** 10.1101/2024.08.28.610181

**Authors:** Biswajoy Ghosh, Komal Agarwal, Anowarul Habib, Krishna Agarwal, Frank Melandsø

## Abstract

3D cell cultures, including spheroids, have become essential tools in cancer research and drug discovery due to their ability to more accurately mimic in-vivo tissue environments compared to traditional 2D cultures. However, imaging these thick, complex structures remains a challenge, as conventional optical microscopy techniques are limited by shallow depth penetration. This study explores the complementary use of gradient light interference microscopy (GLIM) and scanning acoustic microscopy (SAM) for label-free imaging of 3D spheroid clusters embedded in hydrogels. GLIM offers high-resolution optical imaging but struggles with depth in dense samples, while SAM provides greater depth penetration and a larger field of view, albeit with lower resolution. By correlating SAM and GLIM imaging, this study demonstrates how the two techniques can be synergistically used to enhance the visualization of spheroids, capturing both large-scale structural features and fine cellular details. The benefits make such a platform suitable for screening high-number multi-well plates and evaluating necrotic and angiogenic features from the core of the thick sample. Such platforms have the potential of combining acoustic and optical imaging modalities for high-throughput screening and physical characterization in 3D cell culture research, advancing our understanding of drug efficacy in complex biological systems.

## Introduction

In recent years, three-dimensional (3D) cell cultures, including spheroids and organoids, have emerged as pivotal tools in basic biological research, cancer studies, and drug discovery. These 3D models provide a more accurate representation of the in-vivo cellular environment compared to traditional two-dimensional (2D) cell cultures, which often fail to mimic the complex interactions and physical properties of actual tissues^1,2^. As a result, spheroids are increasingly used to evaluate drug toxicity and treatment efficacy, particularly in the development of new cancer therapies^3^.

Despite their advantages, imaging 3D cell cultures presents significant challenges. Optical microscopy, a commonly used technique for high-throughput screening, is limited by its narrow field of view and shallow depth of penetration. In optical imaging, infrared-based imaging has been useful for deep tissue visualization and determining functional parameters^4^. However, they are not ideal for samples like spheroids that do not differentially absorb infrared light. Tomography provides cross-sectional visualization of thick biological samples and provides details of inner structures. While imaging modalities like CT and MRI provide the highest trans-body penetration, they can be toxic or expensive^5–7^. These limitations hinder the comprehensive visualization of spheroids and organoids, which is crucial for detailed biological studies^8^. To observe cellular dynamic changes, fluorescence-based methods such as fluorescence resonance energy transfer (FRET)^9,10^, fluorescence lifetime imaging microscopy (FLIM)^11,12^, and fluorescence recovery after photobleaching (FRAP)^13,14^have been developed. Additionally, super-resolution techniques like structured illumination (SIM)^15,16^, stimulated emission depletion (STED)^17,18^, and single-molecule localization microscopy (SMLM)^19–21^ allow for the observation of minute details with remarkable precision. The increasing demand for rapid, live imaging of thick 3D samples has driven the development of advanced imaging technologies such as multiphoton^22,23^and light-sheet microscopy^24,25^. Multiphoton microscopy can be used for label-free imaging via 2-photon autofluorescence and 3-photon autofluorescence, which are key to detecting metabolic markers (like Flavin adenine dinucleotide (FADH) and nicotinamide adenine dinucleotide (NADH)). Although light sheet and multi-photon are gentle imaging techniques, they are susceptible to high scattering in dense samples, giving a low signal-to-noise ratio.

Multi-harmonic imaging provides label-free imaging from non-centrosymmetric structures (like collagen, myosin etc.) and lipid water interfaces. Advancements in supporting technologies, such as faster cameras, high-throughput adaptive automation, and innovative dyes, have significantly broadened the scope of these techniques in biological research.

However, fluorescence-based methods are inherently limited when it comes to chemically unperturbed evaluation of biological samples, as they rely on the intervention of labels and associated chemicals. Label-free imaging methods such as brightfield, phase contrast, and differential interference contrast (DIC) microscopy are commonly used alternatives. Brightfield microscopy requires staining for contrast, which poses challenges for live cell imaging. Phase contrast and DIC imaging enhance contrast by optically heightening differences between the sample and the background, yet they cannot quantify phase changes accurately due to the non-linear relationship between intensity values and phase information. Recent advancements in label-free imaging have enabled high-resolution and high-speed imaging of morphology, dynamics, functionality, material exchange, pathogen interaction, biochemistry, and biomechanics.

Balancing depth and resolution is a key aspect of any 3D label-free imaging of thick samples. For a long time, quantitative phase microscopy (QPM) struggled with imaging thick samples due to poor handling of multiple scattered lights that washed out details and resulted in poor image quality. However, with the advent of gradient light interference microscopy (GLIM), phase quantification is achievable for several hundreds of microns into the sample, with the ability to create tomograms enabling cross-sectional visualization^8^. GLIM outshines its predecessors in measuring embryo viability, conducting physiological studies of 3D cultures, engineered tissues, organoids, and living organisms (like C. elegans and zebrafish)^26–28^. GLIM is the label-free analog of confocal microscopy in terms of its ability to optically section the sample by suppressing out-of-focus scattered signals, with resolution limited only by the optical diffraction limit.

While GLIM provides a significant advancement in the imaging of thick biological samples, it still does not address the measurement of mechanical properties such as stiffness. The mechanical properties of spheroids, such as stiffness, serve as important markers in biological research, yet these properties are difficult to measure using traditional optical methods. Scanning acoustic microscopy (SAM) offers a promising alternative, providing a larger field of view, greater depth of penetration, and rapid scanning times^29–31^. This technique enables the measurement of physical properties of spheroids in label-free and live conditions, thus overcoming many of the limitations associated with optical microscopy^32,33^. Previously, high-frequency ultrasound monitoring has been used for real-time imaging of thermal therapy in tissues at microscopic resolution^34^.

A crucial component of evaluating drug screening efficacy is to determine the effect of the drug in the deep tissues, complex organoids, and engineered tissues that are relatively thick. In several cases, the efficacy of the drug is directly tied to its ability to penetrate the target specimen. Given that optical imaging methods are limited by penetration depth, SAM presents a valuable alternative for evaluating changes in the deeper regions of biological specimens^29^. The physiological importance of depth imaging of these 3D samples in drug discovery includes regions of necrosis, angiogenesis, etc. Here, we evaluate the advantage of SAM in-depth imaging of 3D spheroids embedded in hydrogels and benchmark with state-of-the-art optical imaging (confocal). We also evaluate the loss of resolution in SAM imaging and the reconciliation required to provide enough evidence of drug efficacy. We highlight SAM’s complementarity along with label-free and fluorescently labelled microscopy techniques. Our findings reveal the potential of SAM to enhance high-throughput screening and provide more detailed insights into the physical characteristics of spheroids, thereby advancing the field of biomedical research.

## Materials and methods

### Sample Preparation

The spheroids were grown by using human cervical adenocarcinoma cell line HeLa. The spheroids were embedded in Gelatin methacryloyl (GelMA) hydrogel to provide a tissue-like environment. The GelMA was made in-house by using the protocols mentioned previously^35^. The tumor spheroids with a mean diameter of (100*±* 30)*µm* were carefully placed using a micropipette in the GelMA solution. The spheroid clusters were made by aggregating individual spheroids such that some were distributed across larger depths or a larger area. To reduce potential backscatter from the bottom surface during the acoustic imaging, the spheroids were placed approximately 200*µm* above the substrate. The hydrogels were solidified by using 395 nm ultraviolet light for 10 seconds in a *µ*-Slide I Luer 3D with one channel and three wells (ibidi). The samples were kept in a culture medium to ensure they were live during the imaging. SAM imaging was carried out first followed by GLIM imaging. The samples were then fixed and labelled for actin (Phalloidin–Atto 647) and nucleus (Sytox green) for confocal fluorescent imaging. The label-free imaging was conducted within 90 minutes to have the least variables in the sample. The experimental design of the spheroid imaging and the 3D rendered view of collocated optical and SAM image of a spheroid is shown in Figure 1

**Figure 1.**
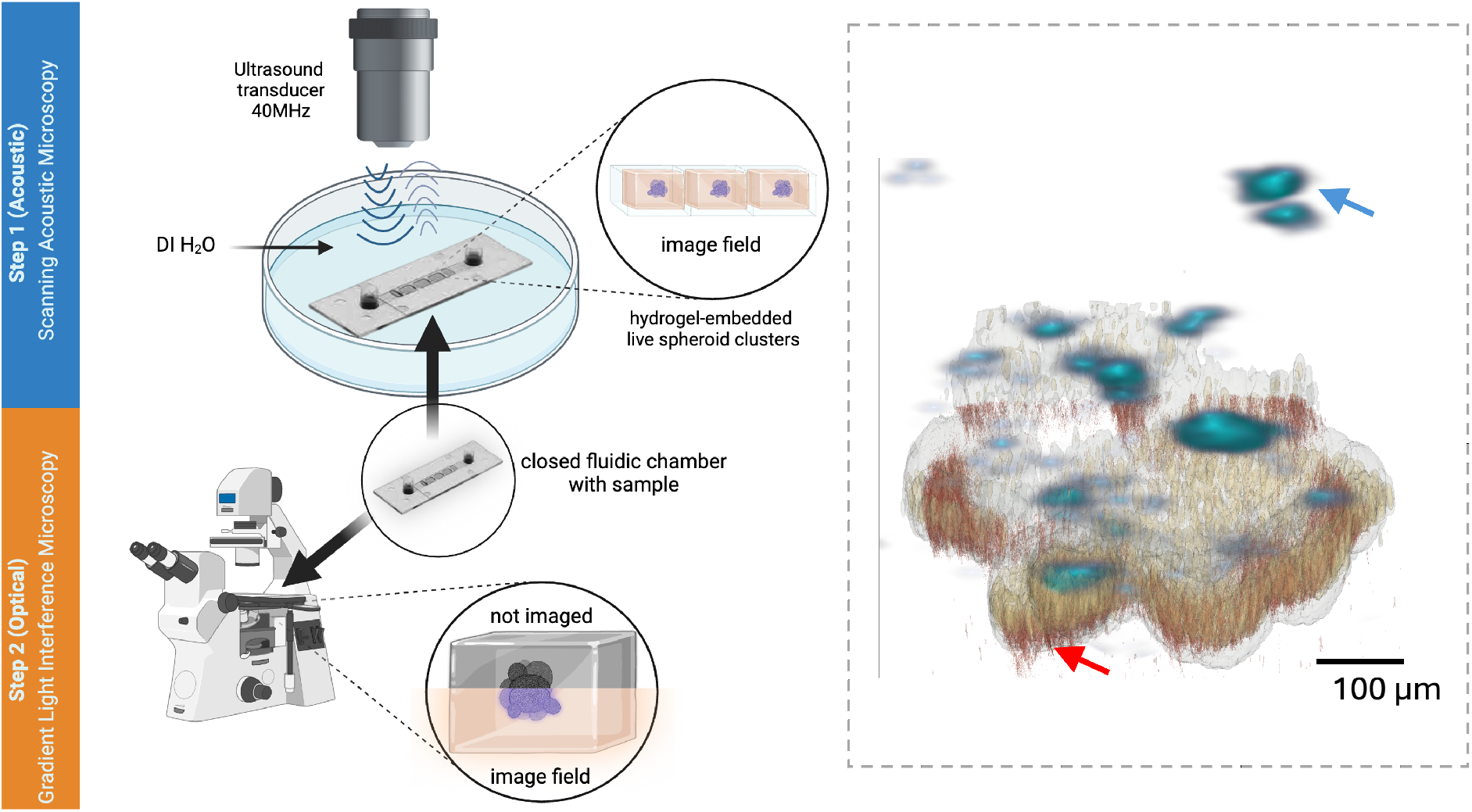
Experimental design: The tumor spheroid clusters were embedded in GelMA hydrogel such that the clusters were spread in all three spatial axes. The sample was imaged with acoustic (SAM) and optical (GLIM) label-free microscopes. SAM of the live sample was performed first, followed by optical imaging with GLIM-an optical imaging modality ideal for thick samples. The sample was later fixed and fluorescently labelled for confocal imaging to compare the results with SAM and GLIM. The study characterizes the imaging outcomes of the two complementary label-free methods for imaging 3D spheroids widely popular for in vitro drug screening applications. SAM provides a larger field of view and deeper penetration, while GLIM provides more details (a higher resolution with a lower field of view and penetration depth). The dotted box on the right displays the 3D-rendered view of the collocated optical (red arrow) and acoustic microscopy, SAM (blue arrow). The image shows superior depth in SAM and better details in optical microscopy.

### Ultrasound Imaging

SAM imaging was performed using a commercially available 40 MHz polyvinylidene fluoride-based polymer spherically focused transducer (Olympus), with an element size of 6.25 mm, an f-number of 2, a focus distance of 12.5 mm, and a -6dB bandwidth. The ultrasonic scanning platform, including the scanning stages, was custom-assembled for the experiments. The setup utilized a Leica DMi8 inverted microscope integrated with an ASI MS-2000 XYZ high-precision scanning stage. Control of the scanning stage and other microscope components was managed using LabVIEW. The ultrasonic functionality was implemented with PXIe FPGA modules and FlexRIO hardware from National Instruments, housed in a PXIe chassis (PXIe-1082). This setup included an arbitrary waveform generator (AT-1212) and an RF amplifier (AMP018032-T) for pulse excitation, along with a 12-bit high-speed (1.6 GS/s) digitizer (NI-5772) for recording reflected signals. Imaging parameters: The images were taken at 25*µm* step size, covering an area 5 × 4 *mm*^2^ at the speed of 1mm/min. The speed of sound and attenuation of sound through the spheroids were used to generate thickness graphs and images of the samples. The penetration depth was up 800*µm*. The observation range of the samples could be adjusted from the data set.

### Optical Imaging

The label-free imaging was done with GLIM (Phi optics) assembled on a Nikon Ti2e inverted microscope. A white transmission light was used for illumination with green filter for live imaging. The confocal imaging for measuring the expression levels was performed with line scanning confocal (Confocal.nl) assembled on the same Nikon microscope. Laser lines of 488nm and 647nm was used for imaging the nucleus and actin. For both GLIM and confocal imaging, we used the objective lens of magnification 20x, 0.45 Numerical Aperture (NA). The z-slice was selected below the Nyquist sampling rate of 1.5um.

## Results

SAM and GLIM imaging of 3D spheroid clusters embedded in hydrogels demonstrate the complementary capabilities of these label-free imaging techniques. Figure 2 shows the complementary aspects and comparisons between these two methods. Figure 2a presents the experimental setup of SAM, and Figure 2b displays the brightfield optical image of the spheroid. SAM enables high-penetration, label-free imaging of live spheroid clusters, as seen in the rendered bottom and side views (Figure 2c-e). The imaging covers a large field of view (310×720×460 µm), providing in-depth visualization of the 3D structure. The color-coded depth map highlights SAM’s capacity to resolve multiple layers within the spheroids, showing its suitability for imaging large, thick biological samples. Complementing SAM, GLIM imaging offers higher resolution with a smaller field of view. The full-field view of GLIM (Figure 2f-g) captures maximum projections using a 20x, 0.45 NA objective lens, effectively resolving individual cells. The 3D-rendered GLIM image (Figure 2g) provides detailed structural information, especially in thinner regions of the sample. However, thicker regions show some data loss. The overlayed images (Figure 2h-k) reveal the complementarity between SAM and GLIM. While SAM provides broader spatial coverage and deeper penetration, GLIM offers enhanced cellular resolution. This combination allows comprehensive 3D imaging, capturing both large-scale structural features and fine cellular details. To establish a ground truth, brightfield and confocal imaging in the same region (Figure 2l-o) confirm the accuracy of spheroid cluster localization and the number of clusters, validating the complementary use of SAM and GLIM for label-free imaging of live spheroids. The confocal images also verify the position and morphology of the spheroid clusters.

**Figure 2.**
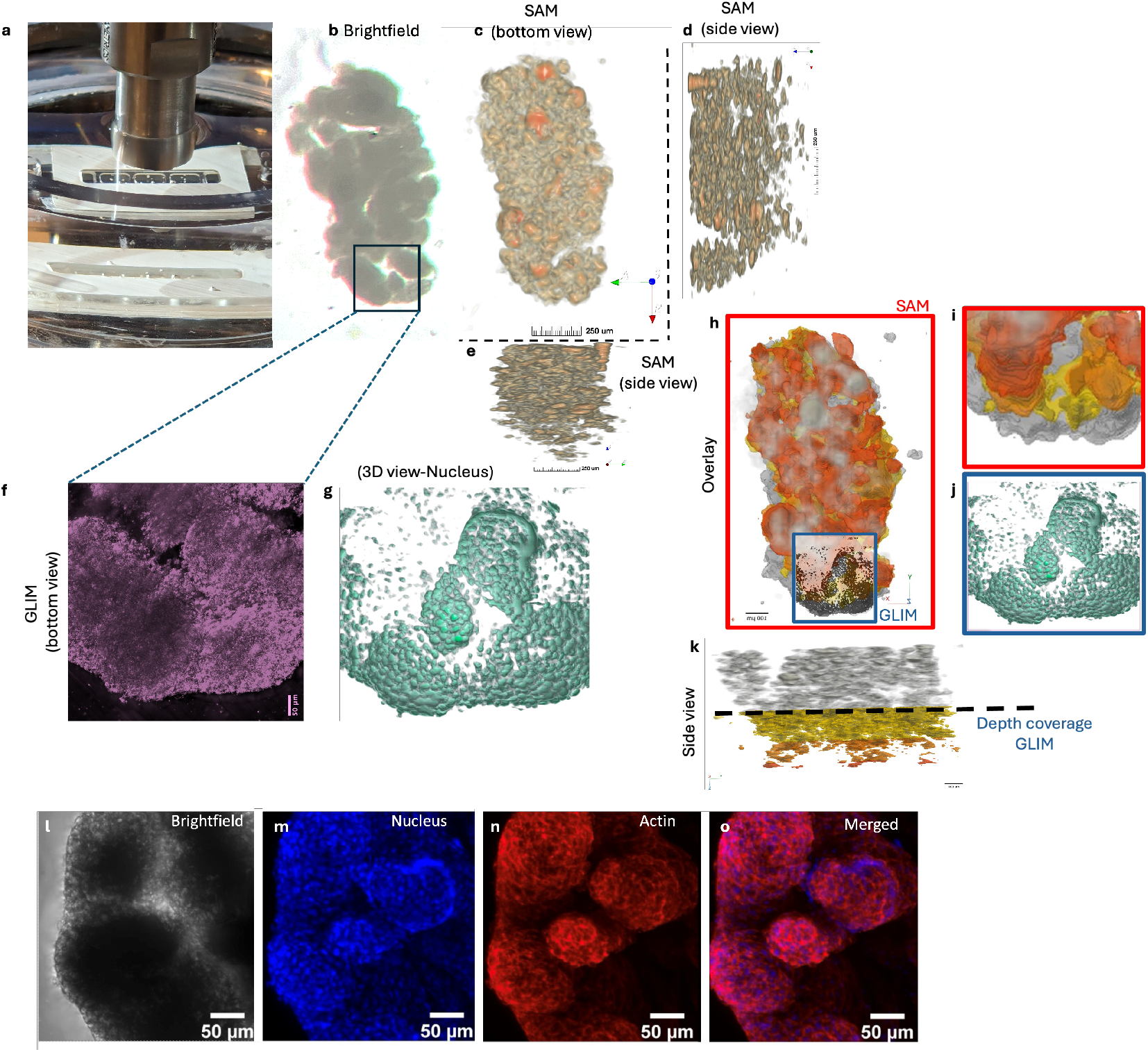
Correlative imaging of 3D tumor spheroid clusters with SAM and GLIM with large area. (a) experimental setup of SAM, (b) a low-resolution brightfield image (transmission) of a spheroid cluster with a large area (xyz 310× 720× 460 *µ*m). (c-e) rendered SAM image showing bottom and side views of the live spheroid cluster, (f) full field view of GLIM image showing max projection using a 20x, 0.45 NA objective lens, (g) 3D rendered GLIM image (h-k) overlayed images of SAM and GLIM showing complementarity in field size and resolution, while SAM shows a much larger field and depth (color-coded depth map), GLIM can resolve individual cells. (l-o) max intensity projection of brightfield and confocal image of the same region, to provide ground truth of number and position of spheroid clusters.

In high-density and thicker spheroid clusters, SAM delineates individual spheroids that are densely packed within 3D space. Ground truth images (Figure 3a-d) reveal a tightly clustered group of spheroids with high z-distribution (dimensions: 31×195×370 µm), with a single spheroid placed to the left for comparison. The SAM imaging setup (Figure 3e) provides etailed bottom and side views (Figure 3f-g), highlighting the distribution of individual spheroid clusters within the dense sample. GLIM images at different z-locations (Figure 3h-k) complement the SAM results by providing additional optical information. However, neither GLIM nor confocal images provide in-depth information about the spheroids, particularly in thicker regions of the sample, due to the limited penetration depth of light. Figure 3l-o shows confocal images of the same region and at the same z-depth locations as GLIM, with overlays of nucleus and actin staining. Confocal staining serves as a reference for comparing optical and acoustic imaging techniques and helps validate the positions and density of the spheroids.

**Figure 3.**
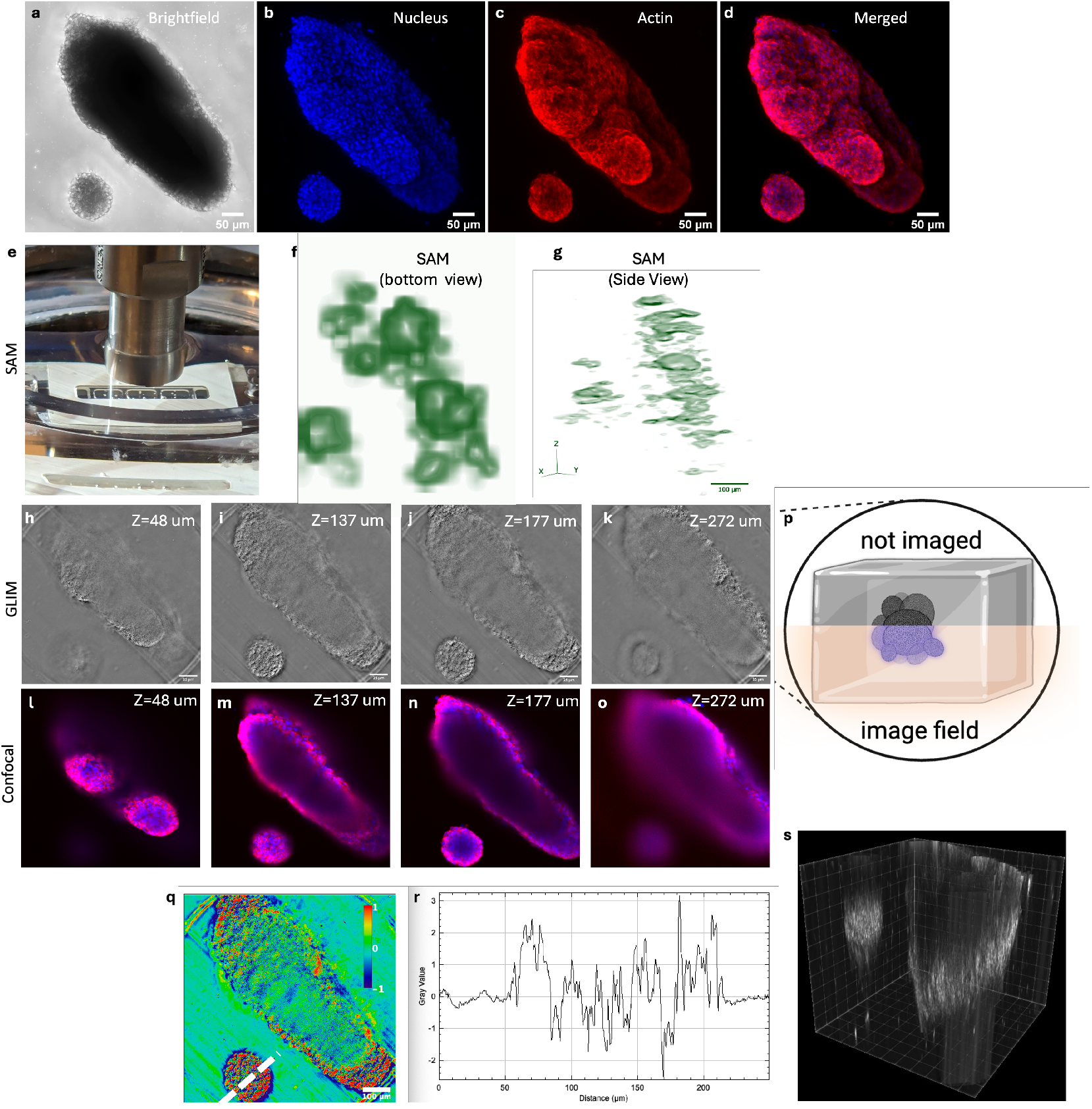
In high-density thicker tumor spheroid clusters, SAM can delineate different spheroids packed densely in 3D space. (a-d) ground truth showing a single spheroid and a densely clustered spheroid with high z-distribution (xyz 110× 415× 370 *µ*m). (e) SAM imaging setup, (f,g) rendered SAM image in bottom and side views to show individual cluster distribution. (h-k) GLIM images of the sample at different z-locations, (l-o) corresponding confocal image showing nucleus (blue) and actin (red). (p) illustrates the imaged and non-imaged fields in the optical image. (q) a quantified phase map to measure the dry mass of the spheroids to detect changes in treatments and stress conditions. (r) a line profile from the single spheroid showing that GLIM can quantify phase information throughout the whole depth of the single spheroid. (s) a 3D rendition of the GLIM image shows that the bottom part of the dense cluster has phase information, while in the upper regions beyond 150 um (farther from the objective lens), no measurable information is derived.

Figure 3p illustrates the imaged and non-imaged fields, highlighting the challenges in capturing all regions optically.Quantitative phase maps (Figure 3q) measure the dry mass and physiological state of cells. A line profile from a single spheroid (Figure 3r) confirms that GLIM can quantify phase information throughout the entire depth of a single spheroid (100 *µm* in size). Additionally, a 3D GLIM rendition (Figure 3s) reveals phase information in the bottom regions of the dense cluster, while regions beyond 150 *µm* from the objective lens lack measurable phase data. These findings demonstrate the combined strengths of SAM and GLIM in imaging high-density spheroid clusters, with SAM providing large-scale structural information and GLIM offering fine-resolution phase data, particularly at shallower depths.

The comparison between acoustic and optical imaging modalities for spheroid cluster imaging is presented in Figure 4. This figure demonstrates the differences in field of view and depth of penetration between the two methods. Figure 4a shows a series of z-slices obtained from SAM reconstructed images, covering a 500*µm* depth of a spheroid cluster. These slices represent the physical top to bottom of the sample, with the uppermost slice corresponding to the top of the spheroid and the lowermost slice representing the deepest part. The SAM images provide decent intra-spheroid resolution structural details throughout the entire depth of the sample, demonstrating superior penetration depth without significant loss in image quality. The red box superimposed on Figure 4a, highlights the limited depth of optical imaging techniques. This range emphasizes the restrictions of optical methods in visualizing the entire sample, particularly when compared to the deeper penetration achieved by SAM. Figure 4b shows brightfield images of the spheroid cluster at different z-slices. These images offer basic structural information but are constrained by the limited optical penetration depth and field of view, as depicted by the red box. Figure 4c displays fluorescence confocal images at corresponding z-slices, with nuclear staining in blue and actin staining in red. Although confocal imaging provides high-resolution visualization of subcellular structures, the depth penetration is restricted, in line with the optical imaging range outlined by the red box. Confocal microscopy is ideal for detailed imaging of superficial layers but cannot penetrate deeper into the sample. Figure 4d presents GLIM images of the spheroid cluster at different depths. Similar to the confocal images, GLIM provides enhanced contrast and structural detail but remains limited in terms of penetration depth, as reflected by the optical range marked in red.

**Figure 4.**
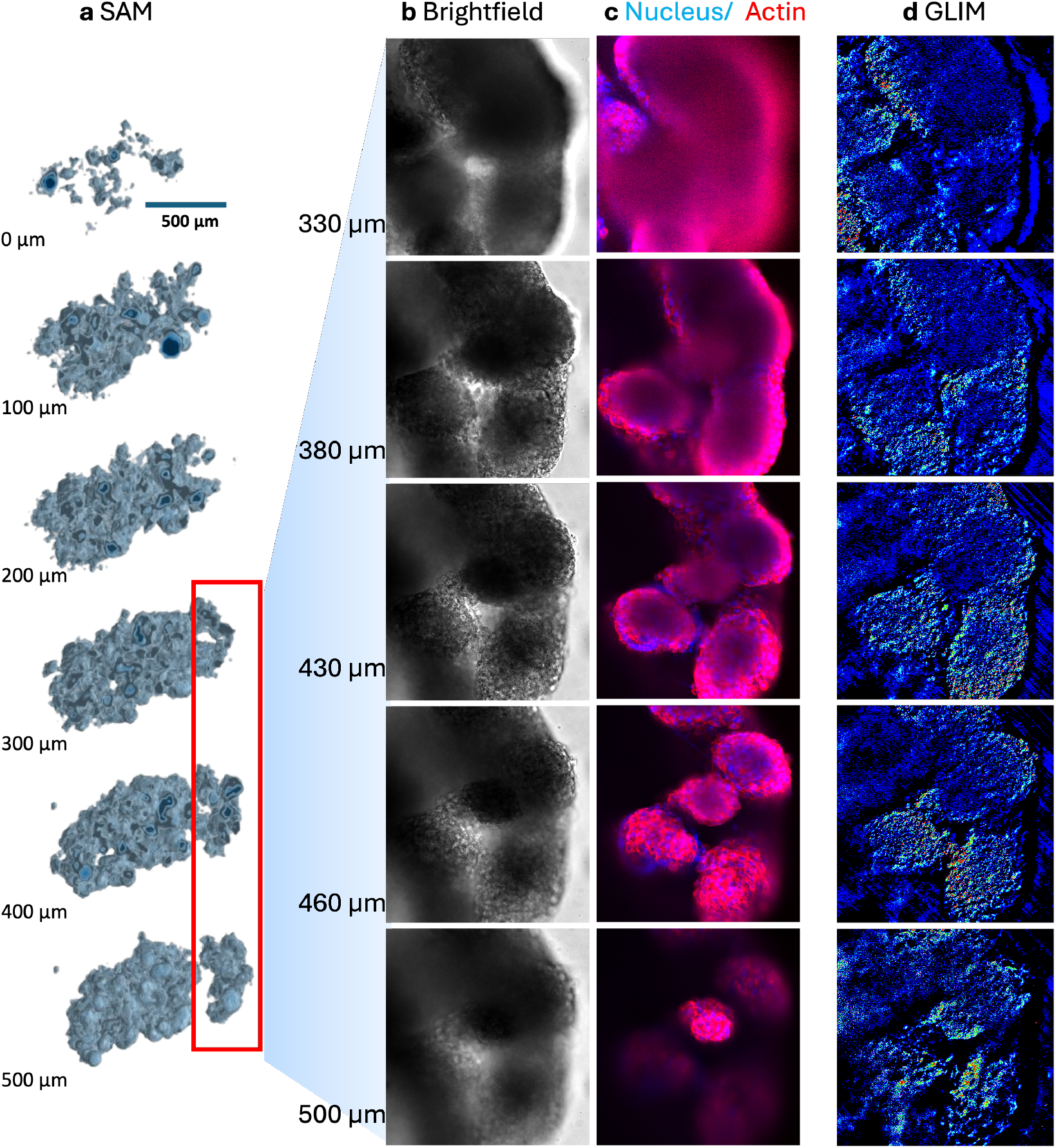
Comparing acoustic and optical field of view and depth of penetration for imaging tumor spheroid cluster. (a) shows the different z-slices of SAM reconstructed image throughout 500 *µ*m depth. The top-to-bottom slices represent the physical top to bottom of the sample. The red box represents the optical imaging range of the sample. The optical counterpart of the spheroid cluster shows (b) brightfield, (c) fluorescence confocal and (d) GLIM image at different z-slices.

Overall, the acoustic imaging modality (SAM) complements optical techniques in terms of depth of penetration, allowing for comprehensive imaging of the entire spheroid cluster. Optical methods, while offering high-resolution images, are restricted to shallower depths, limiting their ability to capture the three-dimensional structure of larger and denser samples fully.

## Discussions and limitations

In the current setup, our objective was to establish the relevance of co-relative SAM-GLIM imaging. We have been able to characterize the benefits and limitations of each modality and the synergy of using the two correlatively. For deep imaging over a larger field of view, which is highly relevant in a drug screening setting, SAM proposes a unique value. Typically, drug screening with spheroids is usually performed in 96-384 well plate formats with the spheroids ranging in sizes between 100*µm* to a few millimeters. The label-free characterization of such samples is crucial to measure drug efficacy in the core of these spheroids which have limited access to nutrition and air availability.

In our study, we preferred to use a multi-spheroids clustered entity instead of a single large spheroid. This choice was intended to use the spatial distribution of the individual spheroids as fiducial markers to correlate the structure between the two modalities. Further, it tests the ability of the microscopes to resolve heterogeneities and asymmetries in the structures.

The acoustic waves interact with the top surface of the sample first and traverse across the entire depth^36,37^. The optical image on the other hand interacts with the bottommost surface first and traverses only halfway across the sample before it attenuates fully. This implies that the bottom half of the sample was the only region that was correlatively imaged. This correlation, therefore, can be subsequently improved.

The GLIM imaging provides an edge when it comes to resolution and quantifying the dry mass of the sample imaged as a function of the gradient phase. The ability of the GLIM for accurate optical sectioning proposes a better lateral and spatial characterization of the sample with diffraction-limited resolution (up to 150 nm). Super-resolution methods can be further employed to reach sub 100*µm* resolution^38–40^. This is relevant for drug screening applications when the efficacy of the lead needs to be determined and connected to the targeted biological pathway.

In bright field images, although the structures are resolvable theoretically, the high density of the sample prohibits the isolation and quantification of relevant biological functions. Confocal imaging shows clear details of target structures but only after the sample is fixed and labelled, which restricts real-time evaluation of biological function. GLIM provides quantitative information on the gradient phase and can optically section individual slices, and thus is a good candidate for high-resolution quantification in high-density and low-depth samples. While SAM provides functional quantification in high-density large depth with slightly compromised resolution^41^. Moreover, from the ultrasound signals it is possible to derive other acoustic properties including impedance, longitudinal velocity, bulk modulus, and stiffness which we have demonstrated previously^32,42,43^. As several diseases including cancers and fibrosis progress through mechanical changes in the samples, this would be an altogether new parameter to be quantified in the bulk drug screening process^44–46^.

The platform’s high sensitivity and label-free nature allow for continuous monitoring of spheroid growth, viability, and response to therapeutic compounds without altering their native state.

## Conclusion

In summary, this study highlights the complementary strengths of gradient light interference microscopy (GLIM) and scanning acoustic microscopy (SAM) for label-free imaging of three-dimensional (3D) spheroid clusters embedded in hydrogels. GLIM provides high-resolution optical imaging, which is particularly useful for detailed visualization of cellular structures at shallow depths. However, its effectiveness diminishes in thicker samples due to limited penetration depth. Conversely, SAM excels in depth penetration and provides a larger field of view, allowing for the comprehensive imaging of larger and denser samples, despite its lower resolution compared to optical techniques. The correlative use of both modalities enables a more complete characterization of spheroids, capturing both fine cellular details and large-scale structural features, making this combined approach highly suitable for high-throughput screening applications in cancer research and drug discovery.

## Acknowledgements

The research is funded by the following projects: Research Council of Norway projects-“Fiber Resolution Targets for Optical Nano and Microscopes (FiRsT)” (349934) and “Cyto-Motility and Cyto-Plasticity in Vitro Live-Cell Assay” (345442); Universitetet i Tromsø (UiT) grants-UiT Talent Supplementary and UiT Talent Innovation Main Project “CYMOPLIVE”, and “VirtualStain” (2061348); H2020 FET-Open RIA project “OrganVision” (964800) and ERC Starting Grant (804233). We also acknowledge UiT The Arctic University of Norway for funding the article processing charges of the published article.

## Author contributions statement

F.M, B.G, Ko.A. conceived the idea and were involved in the experiment. B.G and Ko.A wrote the manuscript. F.M performed ultrasound imaging and analysed the results. B.G performed optical experiments and analyzed the data. A.H helped with characterization of SAM. Kr.A provided resources and infrastructure for cell culture. All authors reviewed the manuscript.

